# Molecular architecture of the SARS-CoV-2 virus

**DOI:** 10.1101/2020.07.08.192104

**Authors:** Hangping Yao, Yutong Song, Yong Chen, Nanping Wu, Jialu Xu, Chujie Sun, Jiaxing Zhang, Tianhao Weng, Zheyuan Zhang, Zhigang Wu, Linfang Cheng, Danrong Shi, Xiangyun Lu, Jianlin Lei, Max Crispin, Yigong Shi, Lanjuan Li, Sai Li

**Affiliations:** State Key Laboratory for Diagnosis and Treatment of Infectious Diseases, Hangzhou 310003, China; National Clinical Research Center for Infectious Diseases, First Affiliated Hospital, Zhejiang University School of Medicine, Hangzhou 310003, China; School of Life Sciences, Tsinghua University, Beijing 100084, China; Beijing Advanced Innovation Center for Structural Biology & Frontier Research Center for Biological Structure, Beijing 100084, China; Tsinghua University-Peking University Joint Center for Life Sciences, Beijing 100084, China; School of Biological Sciences, University of Southampton, Southampton SO17 1BJ, UK; Key Laboratory of Structural Biology of Zhejiang Province, School of Life Sciences, Westlake University, 18 Shilongshan Road, Hangzhou 310024, Zhejiang Province, China; Institute of Biology, Westlake Institute for Advanced Study, 18 Shilongshan Road, Hangzhou 310024, Zhejiang Province, China

## Abstract

Severe acute respiratory syndrome coronavirus 2 (SARS-CoV-2) is an enveloped virus responsible for the COVID-19 pandemic. Despite recent advances in the structural elucidation of SARS-CoV-2 proteins and the complexes of the spike (S) proteins with the cellular receptor ACE2 or neutralizing antibodies, detailed architecture of the intact virus remains to be unveiled. Here we report the molecular assembly of the authentic SARS-CoV-2 virus using cryo-electron tomography (cryo-ET) and subtomogram averaging (STA). Native structures of the S proteins in both pre- and postfusion conformations were determined to average resolutions of 8.7-11 Å. Compositions of the N-linked glycans from the native spikes were analyzed by mass-spectrometry, which revealed highly similar overall processing states of the native glycans to that of the recombinant glycoprotein glycans. The native conformation of the ribonucleoproteins (RNP) and its higher-order assemblies were revealed. Overall, these characterizations have revealed the architecture of the SARS-CoV-2 virus in unprecedented detail, and shed lights on how the virus packs its ∼30 kb long single-segmented RNA in the ∼80 nm diameter lumen.

## INTRODUCTION

As of August 10^th^, 2020, a total of over 20 million cases of COVID-19 were reported and more than 700 thousand lives were claimed globally (https://covid19.who.int). The causative pathogen, SARS-CoV-2, is a novel β-coronavirus (Lu et al., 2020; Wu et al., 2020; Zhou et al., 2020). SARS-CoV-2 encodes at least 29 proteins in its (+) RNA genome, four of which are structural proteins: the spike (S), membrane (M), envelope (E) and nucleocapsid (N) proteins (Kim et al., 2020).

The ∼600 kDa, trimeric S protein, one of the largest known class-I fusion proteins, is heavily glycosylated with 66 N-linked glycans (Walls et al., 2020; Watanabe et al., 2020a; Wrapp et al., 2020). Each S protomer comprises the S1 and S2 subunits, and a single transmembrane (TM) anchor (Wrapp et al., 2020). The S protein binds to the cellular surface receptor angiotensin-converting enzyme-2 (ACE2) through the receptor binding domain (RBD), an essential step for membrane fusion (Hoffmann et al., 2020; Lan et al., 2020; Shang et al., 2020; Wang et al., 2020; Yan et al., 2020; Zhou et al., 2020). The activation of S requires cleavage of S1/S2 by furin-like protease and undergoes the conformational change from prefusion to postfusion (Belouzard et al., 2009; Kirchdoerfer et al., 2018; Simmons et al., 2004; Simmons et al., 2013; Song et al., 2018). Several prefusion conformations have been resolved for the S protein, wherein the three RBDs display distinct orientations, “up” or “down” (Walls et al., 2020; Wrapp et al., 2020). The receptor binding sites expose, only when the RBDs adopt an ‘up’ conformation. The “RBD down”, “one RBD up” and “two-RBD up” conformations have been observed in recombinantly expressed S proteins of SARS-CoV-2 (Henderson et al., 2020; Walls et al., 2020; Wrapp et al., 2020). Upon activation, S follows a classic pathway among class-I fusion proteins (Rey and Lok, 2018): it undergoes dramatic structural rearrangements involving shedding its S1 subunit and inserting the fusion peptide (FP) into the target cell membrane (Cai et al., 2020). Following membrane fusion, S transforms to a needle-shaped postfusion form, having three helixes entwining coaxially (Cai et al., 2020). Despite the efforts in elucidating the SARS-CoV-2 virus host recognition and entry mechanism at near-atomic resolution using recombinant proteins, high-resolution information regarding the *in situ* structures and landscape of the authentic virus is in-demand.

Coronavirus has the largest genome among all RNA viruses. It is enigmatic how the N protein oligomerizes, organizes, and packs the ∼30 kb long single-stranded RNA in the viral lumen. Early negative-staining electron microscopy of coronaviruses showed single-strand helical RNPs with a diameter of ∼15 nm (Caul and Eggleston, 1979). Cryo-ET of SARS-CoV revealed that RNPs organized into lattices underneath the envelope at ∼4-5 nm resolution (Neuman et al., 2006). However, such ultrastructure was not observed in the Mouse hepatitis virus (MHV), the prototypic β-coronavirus (Barcena et al., 2009). No molecular model exists so far for the coronavirus RNP, and little is known about the architecture, assembly and RNA packaging of the RNPs of other (+) RNA viruses.

To address these questions, we combined cryo-ET and STA for the imaging analysis of 2,294 intact virions propagated from an early viral strain (Yao et al., 2020). To our knowledge, this is the largest cryo-ET data set of SARS-CoV-2 virus to date. Here we report the architecture and assembly of the authentic SARS-CoV-2 virus.

## RESULTS AND DISCUSSION

### Molecular landscape of the SARS-CoV-2 virus

SARS-CoV-2 virions (ID: ZJU_5) were collected on January 22^nd^, 2020, from a patient with severe symptoms, and were propagated in Vero cells. The patient was infected during a conference with attendees from Wuhan (Yao et al., 2020). For cryo-EM analysis, the viral sample was fixed by paraformaldehyde, which has minor effects on protein structure at 7-20 Å resolution (Li et al., 2016; Wan et al., 2017). Intact and unconcentrated virions were directly visualized from the supernatant by cryo-EM, showing both ellipsoidal and spherical enveloped particles (Figure S1A), consistent with the observation on virions concentrated by ultracentrifugation through sucrose cushion (Figure 1B). We modelled 2,294 virions as ellipsoids by meshing their lipid envelopes, measuring average diameters of 64.8±11.8 nm, 85.9±9.4 nm and 96.6±11.8 nm for the short, medium and long axis of the envelope respectively. S proteins and RNPs (Figures 1A, S1B, S5A and Video S1) represented the most distinctive features of the virus. A comprehensive analysis has been carried out on the data. In total, 56,832 spikes were manually identified from the virions, approximately 97% of which are in the prefusion conformation, 3% in the postfusion conformation (Method details). An average of 26 prefusion S were found randomly distributed on each virion (Figures 1B-C). The spike copy number per virion is comparable to HIV (Liu et al., 2008), but ∼5 times less than the Lassa virus (LASV) (Li et al., 2016) or ∼10 times less than the Influenza virus (Harris et al., 2006). 18,500 RNPs were manually identified in the viral lumen (Table S1), giving an average of 26 RNPs per virion. However, since the viral lumen is tightly packed with RNPs and electron opaque, the actual number of RNPs per virion was estimated to be 20-30% more, i.e. 30-35 RNPs per virion. Regularly ordered RNP ultrastructures were occasionally observed (Figure S5A), indicating the RNPs could form local assemblies.

**Figure 1.**
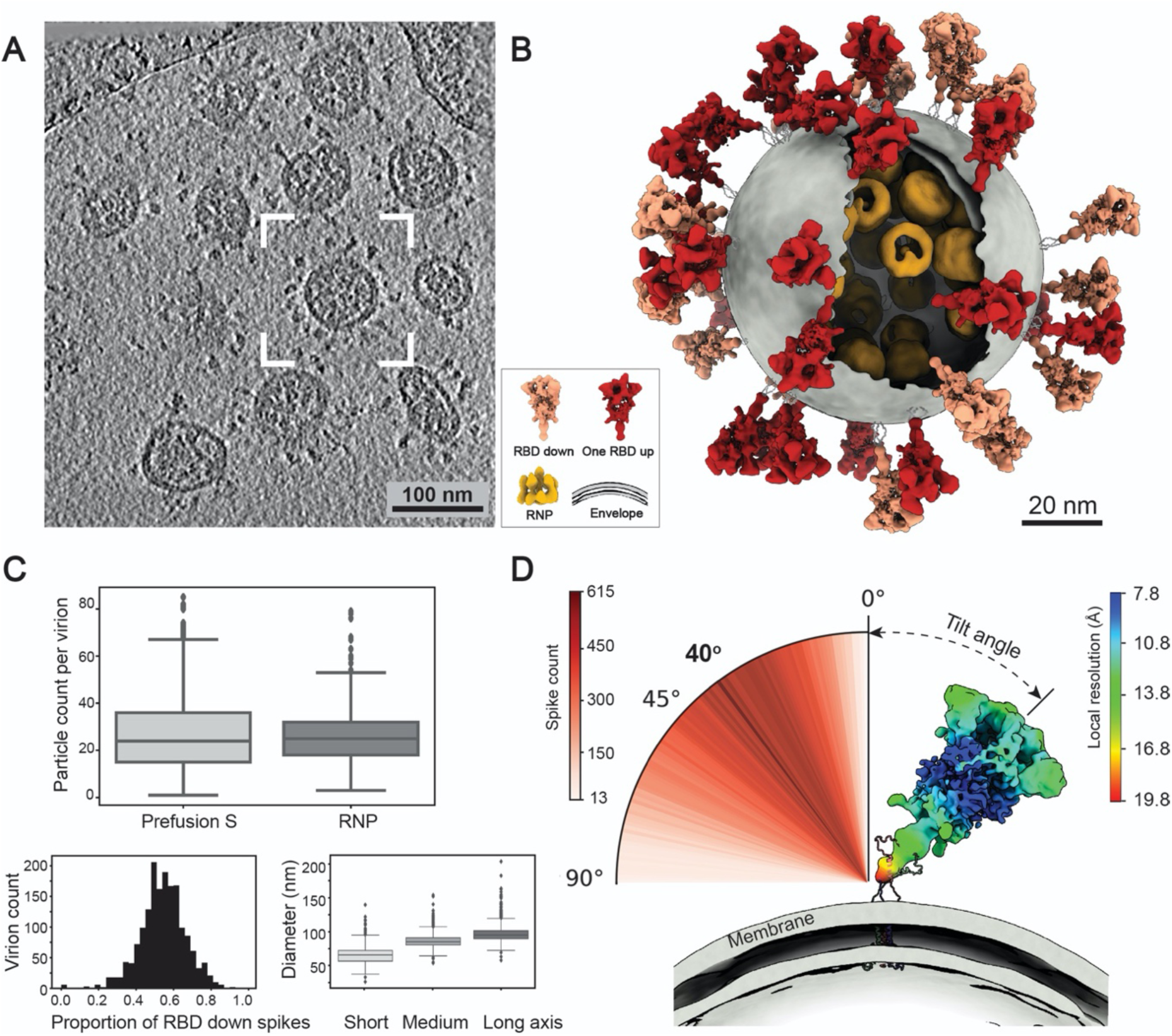
Molecular architecture of the SARS-CoV-2 virus. (A) A representative tomogram slice (5 Å thick) showing pleomorphic SARS-CoV-2 virions. (B) A representative virus (boxed in A) is reconstructed by projecting the prefusion S in RBD down conformation (salmon) and one RBD up conformation (red), lipid envelope (grey), and RNPs (yellow) onto their refined coordinates. RNPs away from the envelope are hidden for clarity. The unsolved stem regions of the spikes are sketched from a predicted model (https://zhanglab.ccmb.med.umich.edu/COVID-19). (C) Summary of structural observations. *Upper*: number of prefusion S and RNPs per virion; *Lower left*: ratio of the S proteins between RBD down and one RBD up conformations; *Lower right*: statistics of the dimension of SARS-CoV-2 viral envelopes. (D) Distribution of the spike tilt angle reveals a prevailing tilt of 40° relative to the normal axis of the envelope. Shown here is a representative RBD down spike in authentic orientation to the envelope (grey). The spike is colored by local resolution, and the predicted model of the stem is fitted for illustration purpose.

Two conformations of the prefusion S, namely the RBD down and one RBD up conformations from inactivated SARS-CoV-2 virions were classified and reconstructed to 8.7 Å and 10.9 Å resolution by STA, with local resolution reaching 7.8 Å (Figures S3A-S3C). The heptad repeat 1 (HR1) and central helix (CH) domains of the S2 subunits represent the best resolved domains (Figure S3D). The proportion of RBD down conformation among all prefusion S was estimated to be 54% per virion (Figure 1C). The membrane proximal stalk of S represented the poorest resolved region with a local resolution of ∼20 Å, showing no trace of the TM or membrane in the structure (Figure 1D). When the tomogram slices were scrutinized, spike populations that either stand perpendicular to or lean towards the envelope were observed (Video S1), suggesting that the TM was averaged out in the map.

Refined orientations of the prefusion S showed they rotate around their stalks almost freely outside the envelope, with the highest possibility of leaning 40° relative to the normal axis of the envelope (Figures 1B and 1D). The rotational freedom of spikes is allowed by its low population density, which is prominently distinct from other enveloped viruses possessing class-I fusion proteins (Harris et al., 2006; Li et al., 2016; Liu et al., 2008). Interestingly, a minor population of Y-shaped spikes pair having two heads and one combined stem were observed (Figures S2A and S2C), which possibly represent spikes intertwined with their stems. These observations suggest that the SARS-CoV-2 spikes possess unusual freedom on the viral envelope. Such unique features may facilitate the virus in exploring the surrounding environment and better engaging with the cellular receptor ACE2, allowing multiple spikes to bind with one ACE2 or one spike with multiple ACE2 simultaneously. However, the sparsely packed spikes on the viral envelope are also more vulnerable to neutralizing antibodies that bind the otherwise less accessible domains (Chi et al., 2020) or glycan holes (Walls et al., 2019). Our observations on the structures and landscape of the intact SARS-CoV-2 are consistent with two other cryo-ET studies appeared at the same period (Turoňová, 2020; Ke et al., 2020).

### Native structures of S in the prefusion conformation

The native structures of S in the RBD down and one RBD up conformations were similar to the rigidly fitted recombinant protein structures (PDB: 6XR8 and 6VYB) (Cai et al., 2020; Walls et al., 2020), except for the N terminus domain (NTD). Comparison between the rigidly and flexibly fitted PDB: 6XR8 suggested that the NTD on the native spike structure shifted 9 Å (centroid distance) away from S2 (Figure S3E). The slight dilation and lower local resolution (Figure S3A and S3B) of the NTD on the native spike against recombinant spike structures was also observed on the other cryo-ET structures (Ke et al., 2020). It is known that NTD exhibits certain mobility as a rigid body (Cai et al., 2020; Walls et al., 2020; Wrapp et al., 2020; Xiong et al., 2020). It is possible that through large date set and classification, the near-atomic resolution cryo-EM reconstruction of the spike represents its metastable conformation, while the cryo-ET reconstructions represent an average of various dynamic states of the NTD.

Ten N-linked glycans are visible on the RBD down and seven on the one RBD up conformations, of which N61, N282, N801, N1098 and N1134 were best resolved. Interestingly, densities for glycans N1158 and N1173/N1194 are visible on the stem of the spike (Figures 2B and S4C). In general, the glycan densities observed on the native spike fit well with the full-length recombinant structure (Cai et al., 2020), however they are bulkier than observed in the TM truncated recombinant spikes (Walls et al., 2020; Wrapp et al., 2020). Similar observations were reported by two recent studies of the native spike structures (Turoňová, 2020; Ke et al., 2020).

**Figure 2.**
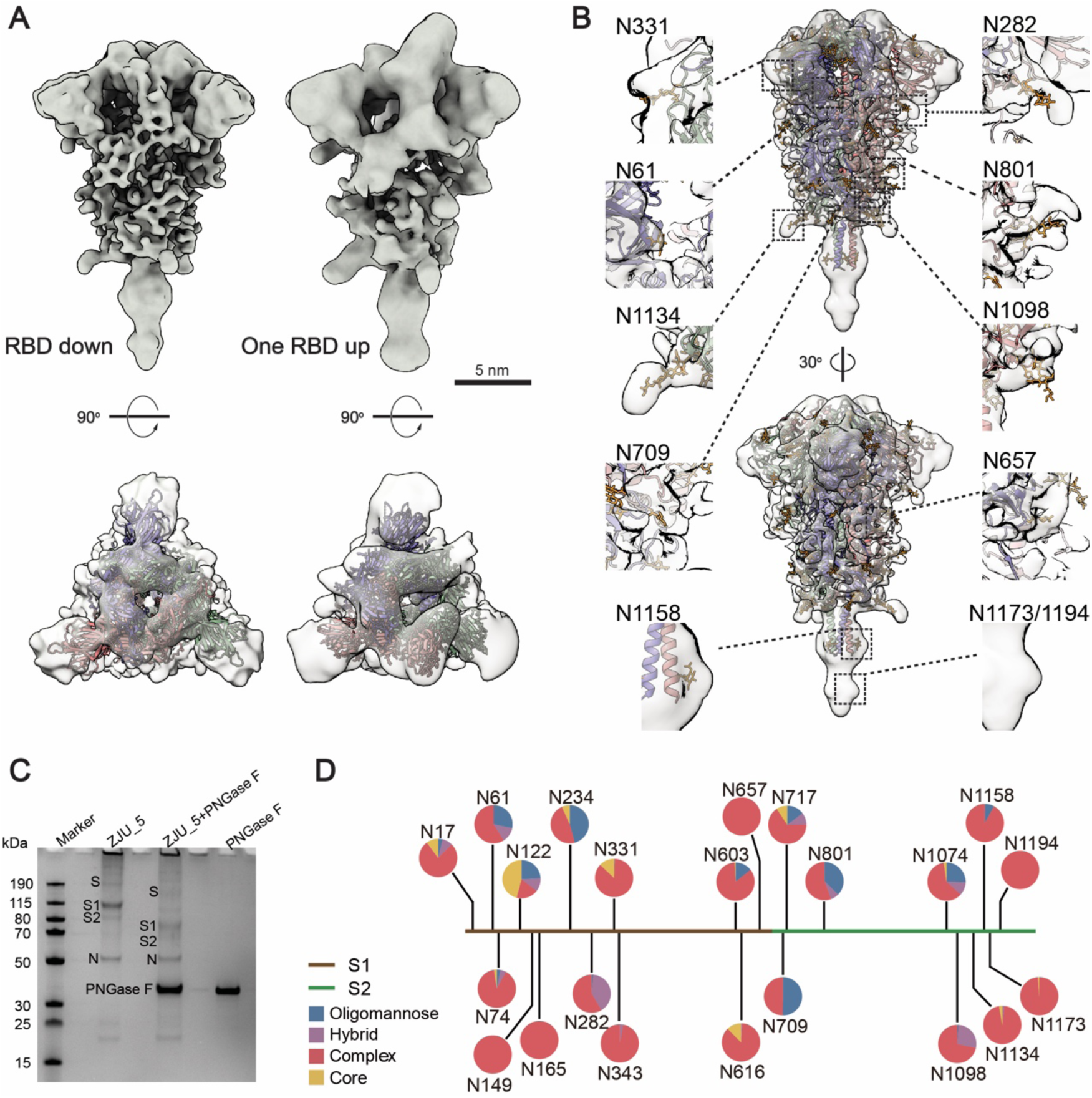
Native structure of the S protein in the prefusion conformation. (A) S in RBD down (resolution 8.7 Å, threshold: 1.2) and one RBD up (resolution 10.9 Å, threshold: 1.5) conformations. Sideview (upper) and topview (lower) of the maps respectively fitted with PDB entries 6XR8 and 6VYB are shown. (B) The RBD down S maps fitted with PDB entry 6XR8. Densities of ten glycans are highlighted in insets. (C) Compositional analysis of the viral sample. The treated viruses, designated ZJU_5, were resolved by SDS-PAGE and visualized by Coomassie blue staining. Lane (1) protein ladder; (2) purified ZJU_5; (3) ZJU_5 treated with PNGase F and (4) PNGase F as control. After PNGase F treatment, the molecular weight of the S1 subunit is reduced by ∼30 kDa, and S2 by ∼20 kDa in weight. (D) Compositional analysis of the surface glycans. The identity and proportion of 22 N-linked glycans from the native S glycans were analyzed by mass spectrometry and presented in pie charts.

We further determined the native glycan identity by analyzing the virus sample using mass spectrometry (MS). The viral particles, with or without PNGase F digestion, were resolved on SDS-PAGE. After PNGase F treatment, the S1 and S2 subunits were reduced by ∼30 kDa and ∼20 kDa, respectively, in weight (Figure 2C). The bands corresponding to S1 and S2 before PNGase F treatment were analyzed by MS to reveal the glycan compositions at each of the 22 glycosylation sites (Figures 2D and S4B). The overall processing states of the native glycans are highly similar to that of the recombinant glycoprotein glycans (Figures S4A and S4B) (Watanabe et al., 2020a), a feature shared with MERS and SARS-CoV (Walls et al., 2019). Populations of under-processed oligomannose-type glycans are found at the same sites as seen in the recombinant material, including at N234 where the glycan is suggested to have a structural role (Casalino et al., 2020). However, as is observed at many sites on HIV (Cao et al., 2018; Struwe et al., 2018), the virus exhibits somewhat lower levels of oligomannose-type glycosylation compared to the recombinant, soluble mimetic. Overall, the presence of substantial population of complex-type glycosylation suggests that the budding route of SARS-CoV-2 into the lumen of endoplasmic reticulum-Golgi intermediate compartments (ERGIC) is not an impediment to glycan maturation and is consistent with both analysis of SARS-CoV glycans (Ritchie et al., 2010) and the identification of neutralizing antibodies targeting the fucose at the N343 glycan on SARS-CoV-2 (Pinto et al., 2020). Furthermore, the lower levels of oligomannose-type glycans compared to HIV and LASV are also consistent with lower glycan density (Watanabe et al., 2020b; Watanabe et al., 2018). Comparing the structures between our native prefusion S to the recombinant ones, we conclude that 1) the N-linked glycans present on the native spike are bulkier and contain elevated levels of complex-type glycans; 2) the recapitulation of the main features of native viral glycosylation by soluble, trimeric recombinant S glycoprotein is encouraging for vaccine strategies utilizing recombinant S protein.

### Native structure of S in the postfusion conformation

Apart from the triangular prefusion S, needle-like densities were occasionally observed on the viral envelope (Figure S2B). A 15.3 Å resolution structure was solved and well fitted with a postfusion form of S (PDB: 6XRA), suggesting a postfusion conformation of the S protein. The ectodomain measures 22.5 nm in length, 6 nm in width and stands perpendicular to the viral envelope. Densities of five N-linked glycans (N1098, N1074, N1158, N1173 and N1194) are displayed on the spike (Figure 3D), reminiscent to the recombinant postfusion structures (Cai et al., 2020; Fan et al., 2020). In comparison to the prefusion conformation, the fixation in orientation to the envelope suggests a dramatic conformational reordering of the stem region to achieve the postfusion conformation. The postfusion S were found only on a small fraction of the viruses (18% of all viruses, each carried an average of 5 postfusion S), hindering us from reaching higher resolutions. Interestingly, among postfusion S-carrying virions, the virions possessing less spikes in total tend to have higher proportion of postfusion S (Figure 3C). Statistics on the refined coordinates (Figures 3A-3C) suggested a tendency of the postfusion S to bundle on the viral surface. One such bundle was reconstructed, showing four postfusion S spaced by ∼10 nm on the viral surface (Figure 3B), compared to ∼15 nm average distance between the nearest prefusion S.

**Figure 3.**
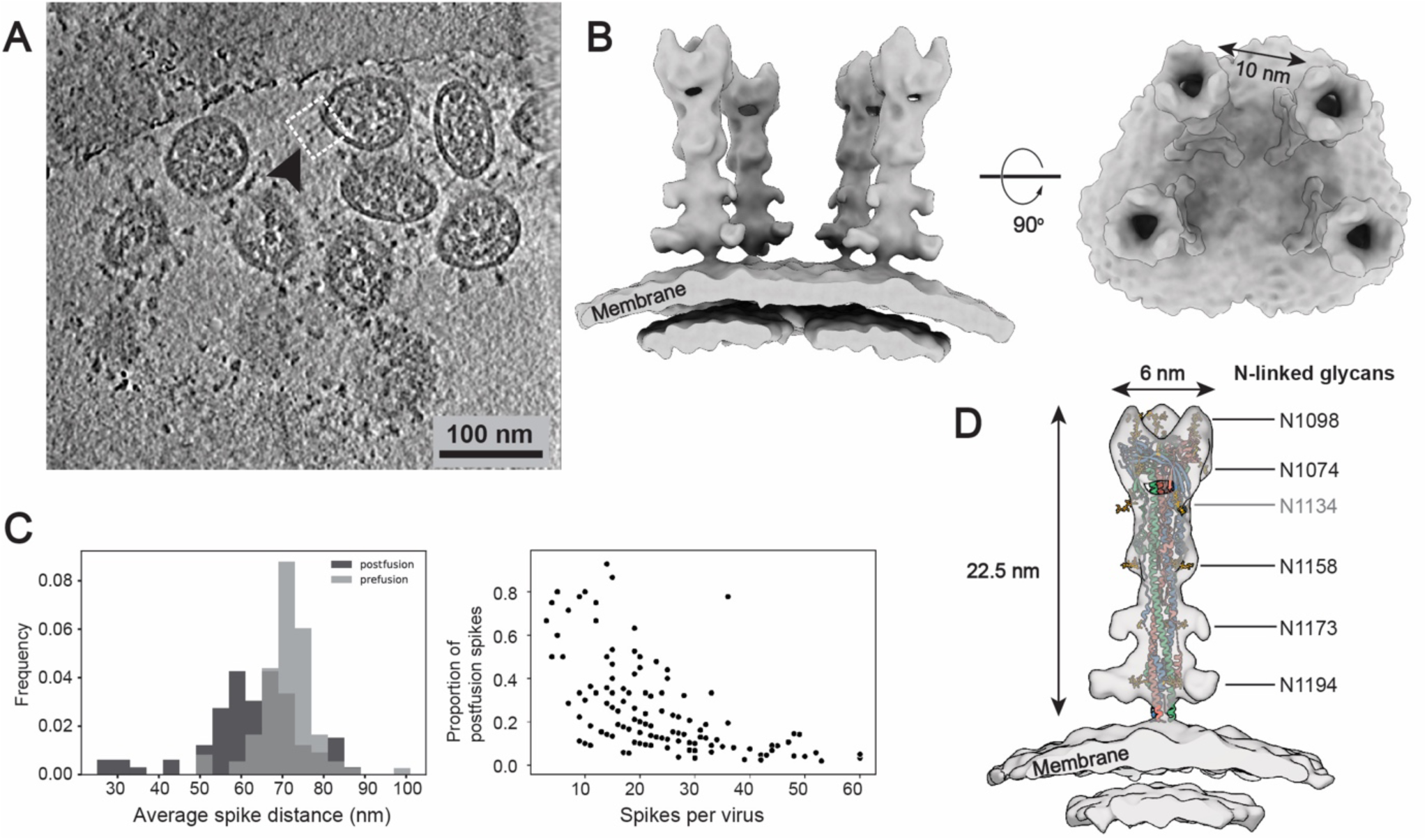
Native structure of S2 in the postfusion conformation. (A) A representative tomogram slice (5 nm thick) showing a cluster of spikes in postfusion conformation on a SARS-CoV-2 virion. (B) 3D reconstructions of S2 in postfusion state. The boxed region in (A) is reconstructed by projecting the solved postfusion S structure onto their refined coordinates. Resolution: 15.3 Å, threshold: 1.0. (C) Distribution of the postfusion S. *Left:* Statistics indicate that the postfusion S are closer to each other on the viral surface compared to the prefusion S; *Right:* viruses possessing less spikes in total tend to have more postfusion S (only virions possessing postfusion S were counted). (D) The postfusion S structure fitted with PDB: 6XRA, displaying densities of N1098, N1074, N1158, N1173 and N1194. Density of N1134 is not visible on the map and is colored grey.

Distinguished from the SARS-CoV virus, which was estimated to possess an average of ∼50-100 spikes per virion (Neuman et al., 2011), the SARS-CoV-2 virus possesses approximately half of the prefusion S and occasionally some postfusion S. The postfusion S observed on the SARS-CoV-2 virus may come from 1) products of occasional, spontaneous dissociation of S1 (Cai et al., 2020), which was cleaved by host proteinases; 2) syncytium naturally formed on infected cells (Xia et al., 2020), when budding progeny virions carried a few residual postfusion S from the cell surface; 3) sample preparation procedure, as cryo-EM images of ß-propiolactone fixed viruses showed most spikes present on the virus are postfusion-like (Liu et al., 2020; Gao et al., 2020). Such instability of the prefusion S was reported on the other β-coronaviruses (Pallesen et al., 2017). In addition, the distribution graph (Figure 3C) implies that the kinetically trapped prefusion S is more fragile than the postfusion S, and could even dissociate from the virus. The speculation is based on the fact that intracellular virions possess more spikes on average (Klein et al., 2020) than the extracellular virions reported by us and the others (Turoňová, 2020; Ke et al., 2020), and the occasional observation of spike-less “bald” virus in our data.

In summary, we speculate that the SARS-CoV-2 prefusion S are unstable, indicating that the distribution of solvent exposed epitopes on the virions is more complicated than the observations on the recombinant proteins. Our observation has implications for efficient vaccine design and neutralizing antibody development, which prefer a sufficient number of stable antigens.

### Architecture and assembly of the RNPs in intact virions

It remains enigmatic how coronaviruses pack the ∼30 kb RNA within the ∼80 nm diameter viral lumen; if the RNPs are ordered relative to each other to avoid RNA entangling, knotting or even damaging; or if the RNPs are involved in virus assembly. When raw tomogram slices were inspected, densely packed, bucket-like densities were discernible throughout the virus lumen, some of which appeared to be locally ordered (Figure S5A). Combining previous cryo-ET observation on coronaviruses (Barcena et al., 2009) and SDS-PAGE/MS analysis (Figure 2C), the densities most likely belonged to the RNPs.

In total 18,500 RNPs were picked in the viral lumen, and initially aligned using a sphere as the template and a large spherical mask. A bucket-like conformation with little structural feature emerged adjacent to the density for lipid bilayer (Figure S5D), suggesting that a significant number of RNPs were membrane proximal. Alignment using a small spherical mask revealed a 13.1 Å resolution reverse G-shaped architecture of the RNP, measuring 15 nm in diameter and 16 nm in height (Figure 4B). Its shape is in good comparison to a recently reported SARS-CoV-2 RNP conformation (Klein et al., 2020), as well as the *in situ* conformation of the chikungunya viral RNP, which is also positive stranded (Jin et al., 2018), but different from the MHV RNP released using detergent (Gui et al., 2017).

**Figure 4.**
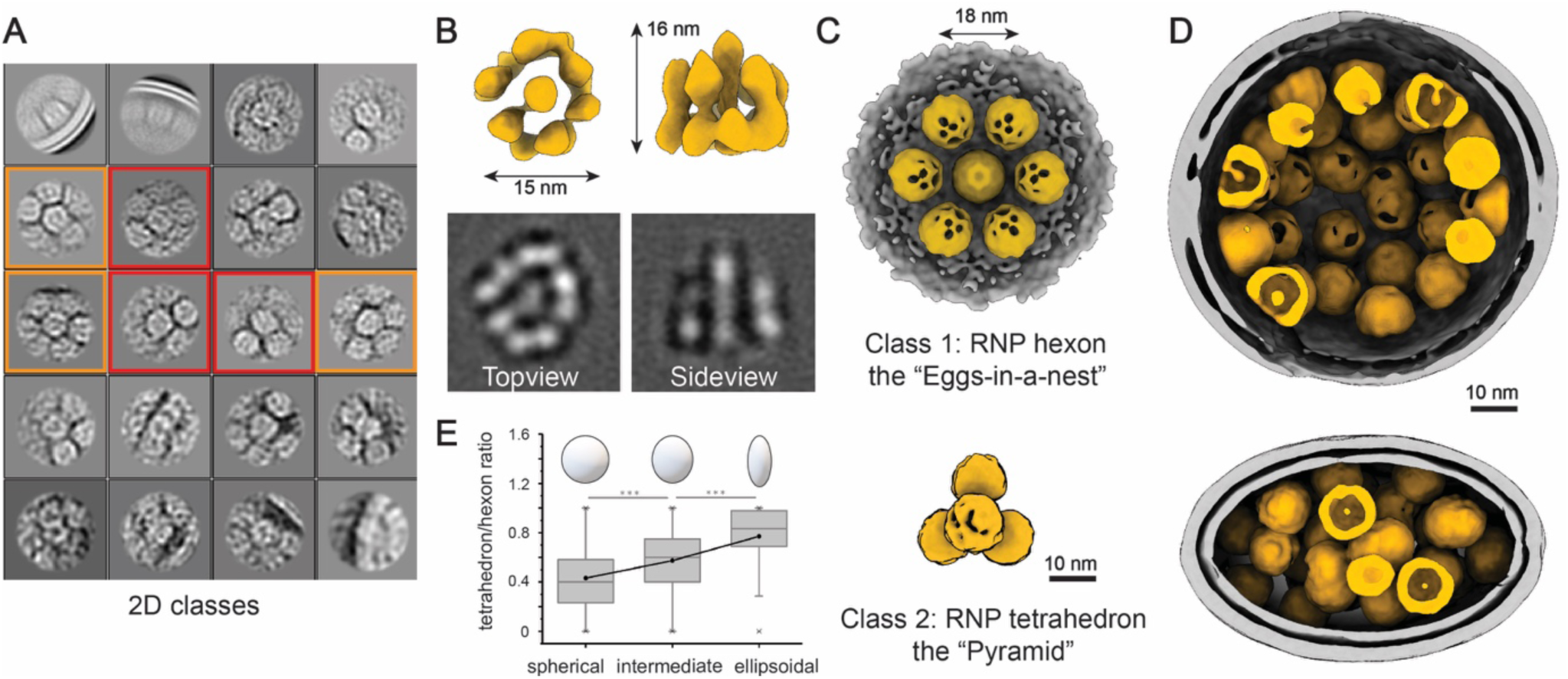
Native assembly of the ribonucleoproteins (RNPs). (A) 2D class averages of RNPs reveal two distinctive types of the RNP ultrastructure: hexameric (boxed in orange) and tetrahedral (boxed in red) assembly. (B) 3D reconstruction and 2D slices of the RNP. Resolution: 13.1 Å, threshold: 1. (C) Ultrastructure of the RNP hexon and tetrahedron assemblies. Seven RNPs are packed against the viral envelope (grey), forming an “eggs-in-a-nest” shaped hexagonal assembly (upper). Four RNPs are packed as a membrane-free tetrahedral assembly (lower), most of which were found inside the virus away from the envelope. Structural features of the RNPs on the assembly are smeared due to the symmetry mis-match between individual RNPs and the assembly. (D) Representative projection of the RNP hexons assembling into a spherical virus, and tetrahedrons into an ellipsoidal virus. (E) Statistics of the ratio of tetrahedron/hexon assembly reveals that the spherical and ellipsoidal virions are more likely to be packed with hexons and tetrahedrons, respectively. The ratio is estimated by sorting 382 virions that have over 5 RNP assemblies by their ratio of long/short axis and counting their ratio of tetrahedron to hexon RNPs.

The map was segmented into five head-to-tail reverse L-shaped densities, each fitted with a pair of N proteins (N terminus domain (N_NTD): 6WKP, C terminus domain (N_CTD): 6WJI) dimerized by the N_CTD (Chen et al., 2007) (Figure S6A and S6C). We analyzed the electrostatic potential distribution on the surface of the decamer, and suggested a tentative structural model of RNA winded RNP (Figures S6B and S6D). Interestingly, an early observation on the MHV RNPs showed ∼15 nm diameter helices with five subunits per turn (Caul and Eggleston, 1979). Due to the limited resolution and little previous structural knowledge about the (+) RNA virus RNPs, our model shall be interpreted with caution.

Further 2D classification of the RNPs revealed three classes: 1) closely packed against the envelope, 2) hexagonally and 3) triangularly packed RNPs (Figure 4A). Following 3D refinement, a membrane proximal, “eggs-in-a-nest” shaped RNP assembly (referred to as the “hexon”), and a membrane-free, “pyramid” shaped RNP assembly (referred to as the “tetrahedron”) emerged (Figures 4C and S5E-S5G). Projection of the two class averages back onto their refined coordinates revealed that the majority of hexons came from spherical virions, while more tetrahedrons from ellipsoidal virions (Figure 4D). This was quantified by statistics: ellipsoidal virions tend to pack more RNP tetrahedrons (Figure 4E). Furthermore, the spacing between two neighboring RNPs (∼18 nm) is the same for both the tetrahedrons and hexons, and some tetrahedrons could assemble into hexons when projected onto their *in situ* coordinates (Figure S5B), suggesting that the RNP triangle is a key and basic packing unit throughout the virus. We further propose that the RNPs are involved in coronavirus assembly and help strengthen the virus against environmental and physical challenges, as purified virions remained intact after five cycles of freeze-and-thaw treatment (Figure S7). Such involvement of RNPs in viral assembly was also reported by (Neuman et al., 2006), who showed RNPs form a lattice underneath the envelope; as well as seen in intracellular virions (Steffen Klein et al., 2020). However, it remains unanswered if the ultrastructures of RNPs are assembled by RNA, the transmembrane M or E proteins, the RNP itself, or multiples of the above.

Solving RNPs to subnanometer resolution was hindered by the crowding of RNPs against each other (Figures S5E-S5G). Furthermore, structural features of the RNPs on higher order assemblies smeared, possibly due to the symmetry mis-match between individual RNPs and the assembly (Figures 4C and 4D). No virus with strictly ordered RNPs throughout the lumen was found by projections. We conclude that the native RNPs are highly heterogeneous, densely packed yet locally ordered in the virus, possibly interacting with the RNA in a beads-on-a-string stoichiometry.

From ∼2,300 intact virions and over 300 tilt series, we provide molecular insights into the structures of spikes in the pre- and postfusion conformations, the RNPs and how they assemble on the authentic virus. We also analyzed the detailed glycan compositions of the native spikes. The reconstructed virus map (Figure 1B) consisting the RBD down S, one RBD up S, lipid envelope and RNP components of the authentic SARS-CoV-2 has been deposited in the Electron Microscopy Data Bank under accession codes EMD-30430, which provide a model for full virus molecular dynamic simulation, 3D printing, education or public media in the future.

## Supporting information

supplementary information for yao et al. cryo-ET of sars-cov-2

## ACKNOWLEDGMENTS

S.L. thank Tsinghua University for providing a Start-up fund, the Tsinghua University Branch of China National Center for Protein Sciences (Beijing) for the cryo-EM facility and the computational facility support, and Dr. Fan Yang, Jie Wen, Danyang Li, Yakun Wang, Anbao Jia and Tao Yang for technical support. We thank Dr. Haiteng Deng and Chongchong Zhao in the Proteinomics Facility at Technology Center for Protein Sciences, Tsinghua University, for protein MS analysis. We thank the computational facility support on the cluster of Bio-Computing Platform (Tsinghua University Branch of China National Center for Protein Sciences Beijing). We are in debt to Dr. Hongwei Wang, Dr. Nieng Yan, Dr. Xinquan Wang, Dr. Lingqi Zhang, Dr. Qiang Ding and Dr. Xueming Li for providing critical advices.

## AUTHOR CONTRIBUTIONS

S.L. conceived and supervised the project. H.Y., N.W., T.W., Z.W., L.C., D.S. and X.L. propagated, fixed and verified the fixation of the virus sample. Y.S. and S.L. isolated the viruses, performed biochemical analysis and prepared the cryo-sample for cryo-EM. Y.S., Y.C., S.L., J.Z. and J.L. collected the EM data. Y.S., Y.C., J.Z., J.X., C.S., Z.Z., and S.L. processed the EM data. S.L. and J.X. performed the subtomogram averaging and classification. Y.C., J.X., S.L. and Y.S. analyzed the structures. C.S., and J.X. performed the statistical analysis. M.C., Y.S. and Z.Z. analyzed the glycan data. S.L., Y.Shi., M.C. and L.L. wrote the manuscript. All authors critically revised the manuscript.

## FIGURES

**Video S1:** A representative raw tomogram (inverted density, lowpassed to 80 Å resolution) overlaid with the reconstruction of a SARS-CoV-2 virus. One of its RBD down S is fitted with PDB: 6XR8.

