## supplementary information for yao et al. cryo-ET of sars-cov-2 for "Molecular architecture of the SARS-CoV-2 virus"

### METHOD DETAILS

#### Sample preparation

SARS-CoV-2 virions isolated from the patient's sputum (ID: ZJU_5) (Yao et al., 2020) were propagated in Vero cells (ATCC CCL-81). Sputum was diluted by 5 volumes of Modified Eagle Medium (MEM) complete medium supplemented with 2% fetal bovine serum (FBS), Amphotericin B (100 ng/ml), Penicillin G (200 units/ml), Streptomycin (200 μg/ml) and centrifuged to remove impurities at 3000 rpm for 10 min in room temperature. Finally, the supernatant was collected and filtered through a 0.45 μm filter. 3 ml of filtered supernatant was added to Vero cells in a T25 culture flask. After incubation at 35 °C for 2 hours to allow binding, the inoculum was removed and replaced with fresh culture medium. The cells were incubated at 35 °C and observed daily to evaluate cytopathic effects (CPE). The SARS-CoV-2 was tested by qRT-PCR and sequencing. For the preparation of enough virus samples, viruses were proliferated using Vero cells in T75 culture flasks. On four days post-infection, 100 ml cell supernatant was cleared from cell debris at 4,000 g centrifugation for 30 min and inactivated with paraformaldehyde (PFA; final concentration 3%) for 48 hours at 4 °C. The supernatant was kept at 4 °C afterwards. All experiments involving infectious virus were conducted in approved biosafety level (BSL)-3 laboratory.

Purification, concentration, biochemical analysis and sample preparation for electron microscopy of inactivated virions were carried out in a BSL-2 lab. For cryo-ET, the fixed virions were pelleted through 30% sucrose cushion by ultracentrifugation (Beckman, IN) at 100,000 g for 3 hours at 4 °C and resuspended in 60 μl HEPES-saline buffer containing 10 mM HEPES, pH 7.3, 150 mM NaCl at 4 °C overnight (Neuman et al., 2008).

#### Deglycosylation and SDS-PAGE

To deglycosylate S protein of virions, 20 μg purified virion sample was treated with 500 units PNGase F (New England Biolabs, Ipswich, MA) at 37 °C for 1 hour. 20 μg of treated and untreated samples were kept at 100 °C for 15 min. Deglycosylation of S protein was characterized by 4 to 12% NuPAGE™ Bis-Tris gel (Invitrogen,Carlsbad, CA). The protein band was stained by One-Step Blue Protein Gel Stain (Biotium, Fremont, CA).

#### Mass spectrometric analysis

For protein identification, the gel bands were excised from the gel, reduced with 5 mM of DTT and alkylated with 11 mM iodoacetamide which was followed by in-gel digestion with sequencing grade modified trypsin (Promega, Madison, WI) in 50 mM ammonium bicarbonate at 37 °C overnight. The sample was quenched by adding 10% trifluoroacetic acid (TFA) to adjust the pH to below 2. The peptides were extracted twice with 0.1% TFA in 50% acetonitrile aqueous solution for 1 hour and then dried in a speedVac. Peptides were dissolved in 25 µl 0.1% trifluoroacetic acid 6 µl of the extracted peptides was analyzed by Orbitrap Fusion Lumos mass spectrometer (Thermo Fisher Scientific, Bremen, Germany).

MS/MS spectra from each LC-MS/MS run were searched against the SARS-CoV-2 database using Proteome Discoverer (Version 1.4) searching algorithm. The search criteria were as follows: full tryptic specificity was required; two missed cleavages were allowed; carbamidomethylation was set as fixed modification; oxidation (M) were set as variable modifications; precursor ion mass tolerance was 20 ppm for all MS acquired in the Orbitrap mass analyzer; and fragment ion mass tolerance was 0.02 Da for all MS2 spectra acquired in the Orbitrap. High confidence score filter (FDR < 1%) was used to select the "hit" peptides and their corresponding MS/MS spectra were manually inspected

For glycan analysis, the gel bands corresponding to S protein were treated with trypsin (Promega, Madison, WI), chymotrypsin (Promega, Madison, WI) and alpha lytic protease (Sigma-Aldrich, St. Louis, MO) similarly to the above procedures.The extracted peptides was analyzed by Orbitrap Fusion Lumos mass spectrometer (Thermo Fisher Scientific, Bremen, Germany).

For LC-MS/MS analysis, the peptides were separated by a 40 min gradient elution at a flow rate 0.30 µl/min with a Thermo-Dionex Ultimate 3000 HPLC system, which was directly interfaced with an Orbitrap Fusion Lumos mass spectrometer (Thermo Fisher Scientific, Bremen, Germany). The analytical column was a home-made fused silica capillary column (75 µm ID, 150 mm length; Upchurch, Oak Harbor, WA) packed with C-18 resin (300 Å, 5 µm, Varian, Lexington, MA). Mobile phase A consisted of 0.1% formic acid, and mobile phase B consisted of 100% acetonitrile and 0.1% formic acid. An LTQ-Orbitrap mass spectrometer was operated in the data-dependent acquisition mode using Xcalibur 4.3.73.11 software and there was a single full-scan mass spectrum in the Orbitrap (300–1500 m/z, 120,000 resolution) followed by 3 s data-dependent MS/MS scans in an Ion Routing Multipole at stepped 27, 30, 33 normalized collision energy (HCD).

Glycopeptide fragmentation data were extracted from the raw file using Byonic™ (Version 2.8.2). The MS data was searched using the Protein Metrics 309 N-glycan library. The search criteria were as follows: Non-specificity; carbamidomethylation (C) was set as the fixed modifications; the oxidation (M) was set as the variable modification; precursor ion mass tolerances were set at 20 ppm for all MS acquired in an orbitrap mass analyzer; and the fragment ion mass tolerances were set at 0.02 Da for all MS2 spectra acquired.

The intensities of each glycan type in identical site were combined and analyzed for proportion. Data with score less than 30 were discarded. The glycans were classfied into oligomannose, hybrid and complex type based on composition. Hybrid and complex type glycan were subdivided according to fucose component and antenna.

**Cryo-electron tomography and electron microscopy**

7 µl virus sample was applied onto a glow discharged copper grid coated with holey carbon (R 2/2; Quantifoil, Jena, Germany), and subsequently dipped onto 500 µl HEPES-saline buffer for 1 second to clear the sucrose. A drop of 3 µl gold fiducial beads (10 nm diameter; Aurion, The Netherlands) was applied and the grid was blotted for 4.5 s, vitrified by plunge-freezing into liquid ethane using a Cryo-plunger 3 (Gatan, CA). Fixed cells cultured on grids were applied with 2 µl fiducial beads (10 nm diameter; Aurion, The Netherlands) prior to single-sided plunge-frozen.

The grids were imaged on a Titan Krios microscope (Thermo Fisher Scientific, Hillsboro, OR) operated at a voltage of 300 kV equipped with an energy filter (slit width 20 eV; GIF Quantum LS, Gatan, CA) and K3 direct electron detector (Gatan, CA). Virions were recorded in super-resolution mode at a nominal magnification of 64,000×, resulting in a calibrated pixel size of 0.68 Å. 361 sets of tilt-series data were collected using the dose-symmetric scheme (Hagen et al., 2017) from -60° to 60° at 3° steps and at various defocus between -1.7 and -5 µm in SerialEM (Mastronarde, 2005). At each tilt, a movie consisting of 8 frames was recorded with 0.0265 s/frame exposure, giving a total dose of 131.2 $e^{-}/Å^{2}$ per tilt series.

For the freeze-and-thaw test, 1.5 µl purified virions were diluted in 6 µl HEPES-saline buffer at 4 °C, then subjected to five cycles of freezing in liquid nitrogen and thawing in water bath at 37 °C. For the negative staining microscopy, 4 µl freeze-and-thaw sample was applied on copper grids (Zhongjingkeyi Technology, Beijing, China), stained using 2% Uranium acetate and imaged using a Tecnai Spirit TEM (Thermo Fisher Scientific, Hillsboro, OR).

**Data processing**

Tilt series data was analysed in a high-throughput pre-processing suite developed in our lab. The electron beam induced motion was corrected using a combination of MotionCor (Li et al., 2013) and MotionCor2 (Zheng et al., 2017) by averaging eight frames for each tilt. Defocuses of the tilt series were measured using Gctf (Zhang, 2016). The tilt series were contrast transfer function corrected using Novactf (Turonova et al., 2017), 319 tilt-series with good fiducial alignment and relative thin ice thickness were reconstructed to tomograms by weighted back projection in IMOD (Kremer et al., 1996), resulting in a final pixel size of 1.36 Å/pixel. The tomograms were 2 × and 4 × binned for subsequent processing. 2,294 virions, 54,878 prefusion S, 2,010 postfusion S and 18,500 RNPs were manually picked. The viral envelopes were manually selected and modelled as ellipsoids using the mesh methods in Dynamo. On average, 425 equally spaced vectors normal to the viral envelope were established per virus. Initial orientations of all spikes were applied using these vectors. The metadata containing all viral parts was organized with Dynamo catalogue (Castano-Diez et al., 2016) for further analysis.

Subtomogram averaging was done using Dynamo (Castano-Diez et al., 2012). For the prefusion S reconstruction, 54,878 subtomograms were extracted from 4 × binned tomograms into boxes of 96×96×96 voxels and EMD-21452 (Walls et al., 2020) was used as the template for their alignment. The resolution was restricted to 40 Å and C3 symmetry was applied at this stage. 8,562 spikes present at the edges of the tomograms were removed to minimize the impact of air-water interface effect and incomplete signal on the structure. The remaining particles were subjected to multi-reference alignment imposing C1 symmetry using EMD-21452 and EMD-21457 lowpassed to 30 Å resolution as the templates, resulting in 25,236 spikes (54.5%) classified into RBD down conformation and 21,080 spikes (45.5%) into one RBD up conformation. Coordinates of the two spike conformations were used to extract boxes of 160×160×160 voxels from the 2 × binned tomograms for further alignment. To prevent overfitting, a customized 'gold-standard adaptive bandpass filter' method was used for the alignment at this stage, and a criterion of 0.143 for the Fourier shell correlation were used to estimate the resolution. The 2 × binned spikes in the RBD down and one RBD up conformations were independently further aligned imposing C3 or C1 symmetry respectively, to 9.5 and 10.9 Å resolution. Finally, the RBD down spike subtomograms were extracted from unbined tomograms into boxes of 256×256×256 voxels and aligned to 8.7 Å resolution. The prefusion S maps were lowpassed according to the estimated local resolutions of the reconstructed subunits. Universal empirical B-factors of -1200 and -2000 were applied to sharpen the RBD down and one RBD up spikes, respectively (Wan et al., 2017).

For the postfusion S reconstruction, 2,010 subtomograms were extracted from 4 × binned tomograms into boxes of 96×96×96 voxels, which were averaged to give an initial template for their alignment. The resolution was restricted to 30 Å and C3 symmetry was applied at this stage. Next, the refined coordinates were used to extract 1,954 postfusion S from the 2 × binned tomograms into boxes of 160×160×160 voxels for gold-standard alignment. Subsequent alignment achieved 15.3 Å resolution.

For the RNP reconstruction, 18,500 manually picked RNPs were extracted into subtomograms of 80×80×80 voxels from 4 × binned tomograms and globally aligned using a large sphere (radius 36 pixels) as the template. The resolution was restricted to 40 Å and no symmetry was applied at this stage. Lipid bilayers were visible in the aligned maps, suggesting part of the RNPs are relatively packed with the membrane. The alignment was repeated using a small spherical mask (radius 18 pixels). A reverse "G"-shaped structure appeared after this stage and the refined coordinates were used to extract particles from the 2 × binned tomograms into boxes of 128×128×128 voxels. Gold-standard was applied to align the RNP to a final resolution at 13.1 Å.

To analyze the local pattern of the RNP assembly, the picked RNPs' coordinates were imported into the Relion subtomogram averaging pipeline (Bharat and Scheres, 2016). The RNP particles were extracted and projected into 2D images. Three characteristic patterns of the 2D classification are selected and subjected to 3D initial model generation and 3D classification: 1) closely packed towards the envelope, 2) hexagonally packed and 3) triangularly packed RNPs. Following 3D refinement, the first class converged only on the membrane. The second class aligned into a hexagonally packed, membrane proximal RNP assembly, and the third class aligned into a tetrahedrally packed, membrane-free assembly. Refined coordinates and orientations of the hexagonal particles (2,270 hexons) and tetrahedral particles (3,659 tetrahedrons) were converted and exported for further alignment in Dynamo. For the RNP hexons, 2,270 particles were extracted into subtomograms of 128×128×128 voxels from 4 × binned tomograms. A shell-shaped mask and C6 symmetry were applied during the alignment. For the RNP tetrahedrons, 3,659 particles were extracted into subtomograms of 90×90×90 voxels from 4 × binned tomograms. A spherical mask and C3 symmetry were applied during the alignment. Both assemblies were only aligned using non-gold standard.

**Assembly structure reconstruction**

Three representative SARS-CoV-2 virus (Figures 1B and 4D) and a bundle of postfusion S (Figure 3B) were reconstructed by projecting all spikes and RNPs onto their refined coordinates and merging the structures using Jsubtomo (Huiskonen et al., 2014). For other map-projection to coordinates (Figure 1D, S2C and S5B), the 'dtplot' function in Dynamo was used. UCSF Chimera (Pettersen et al., 2004) and ChimeraX (Goddard et al., 2018) were used for rendering the graphics.

**Fitting**

Atomic models (PDB accession code 6XR8, 6VYB, 6XRA) of the pre- and postfusion S were rigidly fitted to the corresponding densities using the Fit in Map tool (Pettersen et al., 2004).

The RNP map was segmented into five reverse L-shaped units, which can be further ungrouped into 7 segments above and 10 segments on the base. According to the previous Small-angle X-ray scattering (SAXS) (Chang et al., 2009) and cryo-EM (Gui et al., 2017) reports, the N_NTD and N_CTD possibly form a reverse L-shaped unit; the N_CTD dimer was suggested to be an assembly unit of the RNP (Chen et al., 2007). One segment above and two on the base forming a reverse "L" from the best solved region were selected, and were fitted with a N_CTD dimer (6WJI) using the 'fit to segments' tool (Pintilie et al., 2010) in UCSF Chimera. The segment with the best fitting score (0.78 against 0.74 and 0.72) were adopted as the N_CTD. The other two segments were fitted with the N_NTD monomer (6WKP, score 0.92 and 0.90). With the reverse "L"-shaped N_NTD-CTD pair formed, we fitted the rest of RNP with four such units, leaving two upper segments unoccupied. Together, the map was interpreted as a decamer of N.

Molecular dynamic flexible fitting (MDFF) (Trabuco et al., 2009) was applied to improve the fitting of the atomic model to the S in RBD down conformation. PDB: 6XR8 was prepared in VMD (Humphrey et al., 1996) for the MDFF, which was performed in vacuum until convergence using NAMD 2.12 (Phillips et al., 2005) and the CHARMM36 force field (MacKerell et al., 1998). A scaling factor $\zeta = 0.3$ kcal/mol; the secondary structure, domain and symmetry restraints were applied for the simulation.

### QUANTIFICATION AND STATISTICAL ANALYSIS

#### Morphological description of virions (Figure 1C and S1)

1,959 ellipsoidal models for the envelopes were used for virus diameter estimation statistics; For the statistics in Fig S1, membranes of virion were fitted by ellipse, and 113 virions without high-speed centrifugation and 157 virions with high-speed centrifugation were used.

#### Distribution of spikes in different conformation (Figure 1C and 3C)

1,743 virions were used for calculating the proportion of RBD down/one RBD up conformations per virus; the tilt angle statistics was performed on 39,112 4 × binned spikes (CC > 0.1); and 742 virions were used for RNP statistics. 422 virions were used for the postfusion S statistics. 139 virions which had both prefusion and postfusion S were used for the statistics of conformation proportion. 91 virions which have more than 3 postfusion S and prefusion S were used for spike average distance statistics. And Paired T-test showed average distance among postfusion S and prefusion S is significantly different (p-value<0.01). Statistics was performed using Python package Scipy.

#### Distribution of spikes' tilt angles (Figure 1D)

The aligned location and orientation were used for the statistics of the spike tilt angles. For a given spike on a given virus, the nearest envelope mesh to the spike stem end was found. The angle between vector A (the spike's Z axis) and vector B (normal to the envelope mesh) was calculated. For the statistics in Fig S1C, envelopes of 113 virions before ultracentrifugation and 157 after ultracentrifugation were fitted by ellipse to estimate their diameters.

382 virions with more than 5 tetrahedron/hexon RNP assemblies were included for the statistics shown in Figure 4E. They were sorted into three classes according to the ratio of the longest to the shortest axis: spherical (long/short axis: 1-1.24, 127 virions), intermediate (long/short axis: 1.24-1.43, 127 virions), and ellipsoidal (long/short axis: 1.43-2.45, 128 virions). For each class, the ratio of tetrahedrons to hexons was calculated. Wilcoxon test was applied and showed that the frequency of hexons and tetrahedrons is significantly different between these three classes (p<0.01).

**Supplemental Figures**

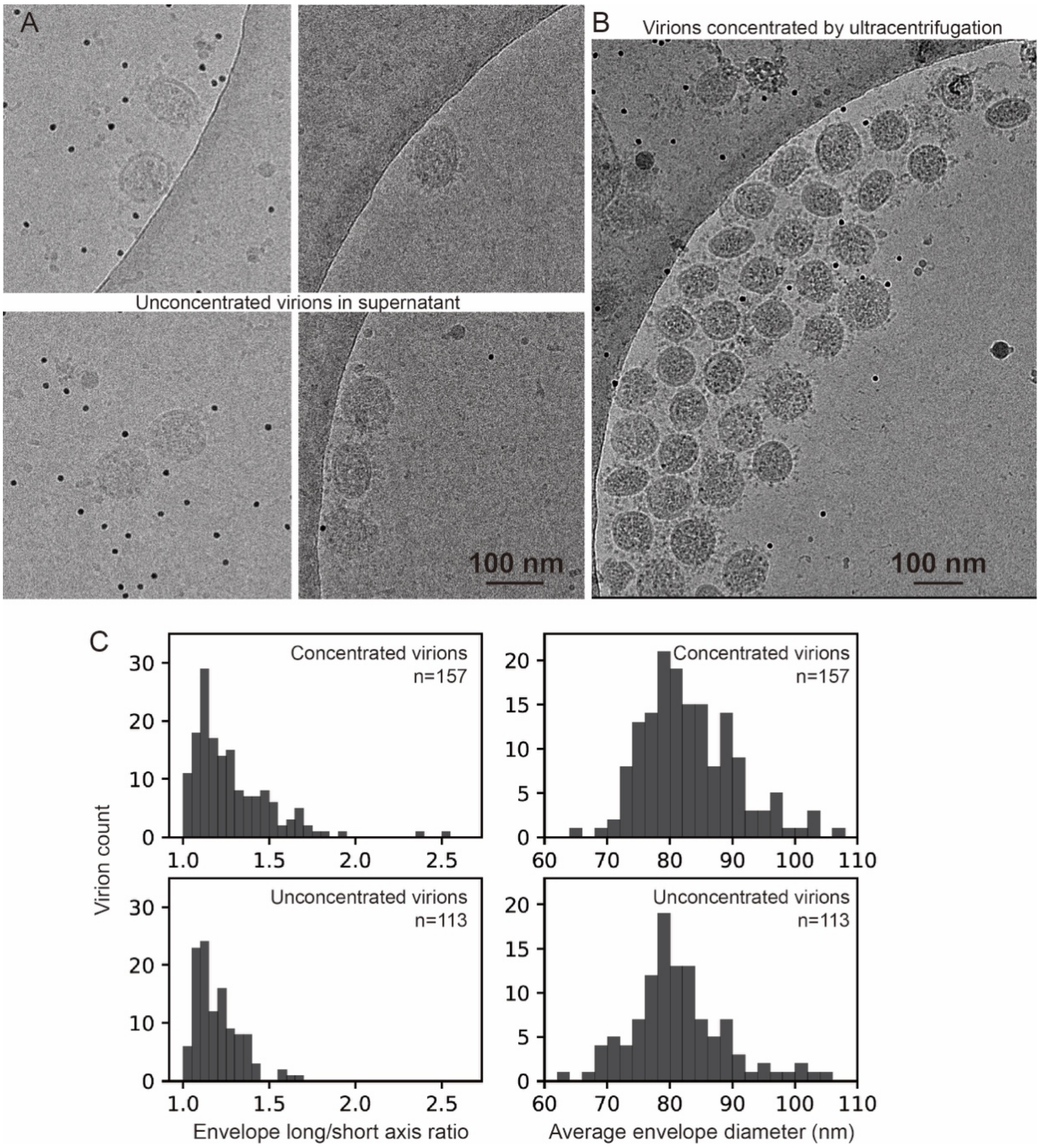


**Figure S1.** Comparison between the unconcentrated and concentrated SARS-CoV-2 virions. Cryo-electron microscopy of virions present in the supernatant of infected Vero cells (A), and virions concentrated by ultracentrifugation (B), showing both spherical and ellipsoidal particles. (C) The long/short axis ratio, as well as the average diameters of the viral envelope measured from the micrographs, are similar between the concentrated (157 virions) and unconcentrated (113 virions) virions.

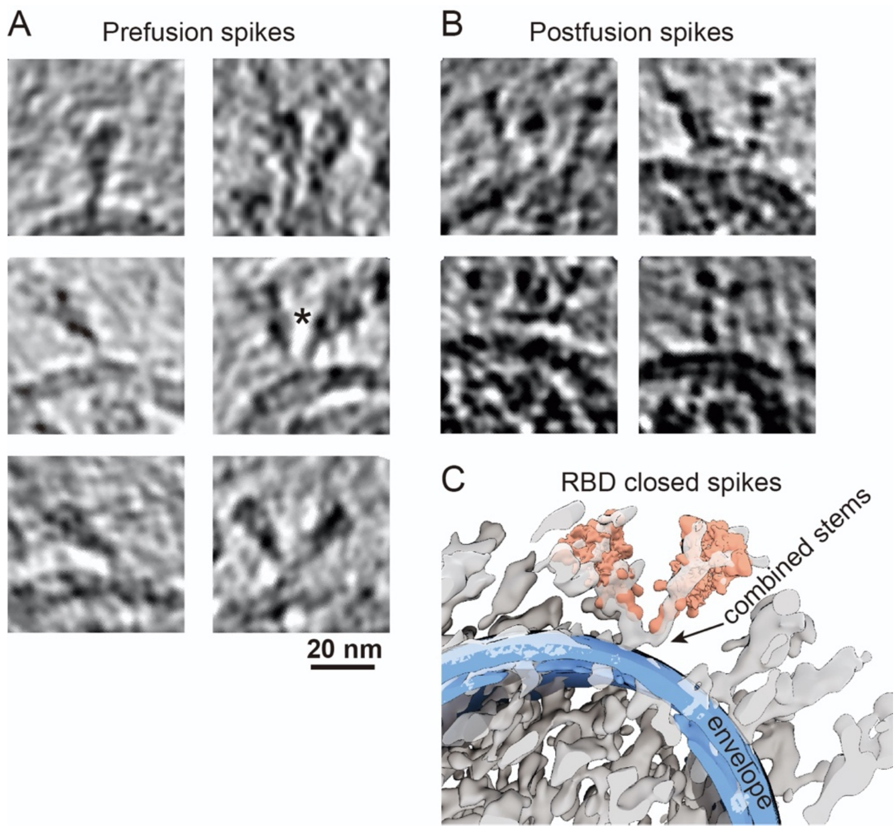


**Figure S2.** Example tomogram slices of spikes in prefusion (5 Å thick) (A) and postfusion (5 nm thick) (B) conformations. (C) A representative pair of Y-shaped spikes in RBD down conformation (marked with * in A) is illustrated by projecting the refined structures onto their coordinates and overlaying with the raw tomogram (lowpassed to 80 Å resolution).

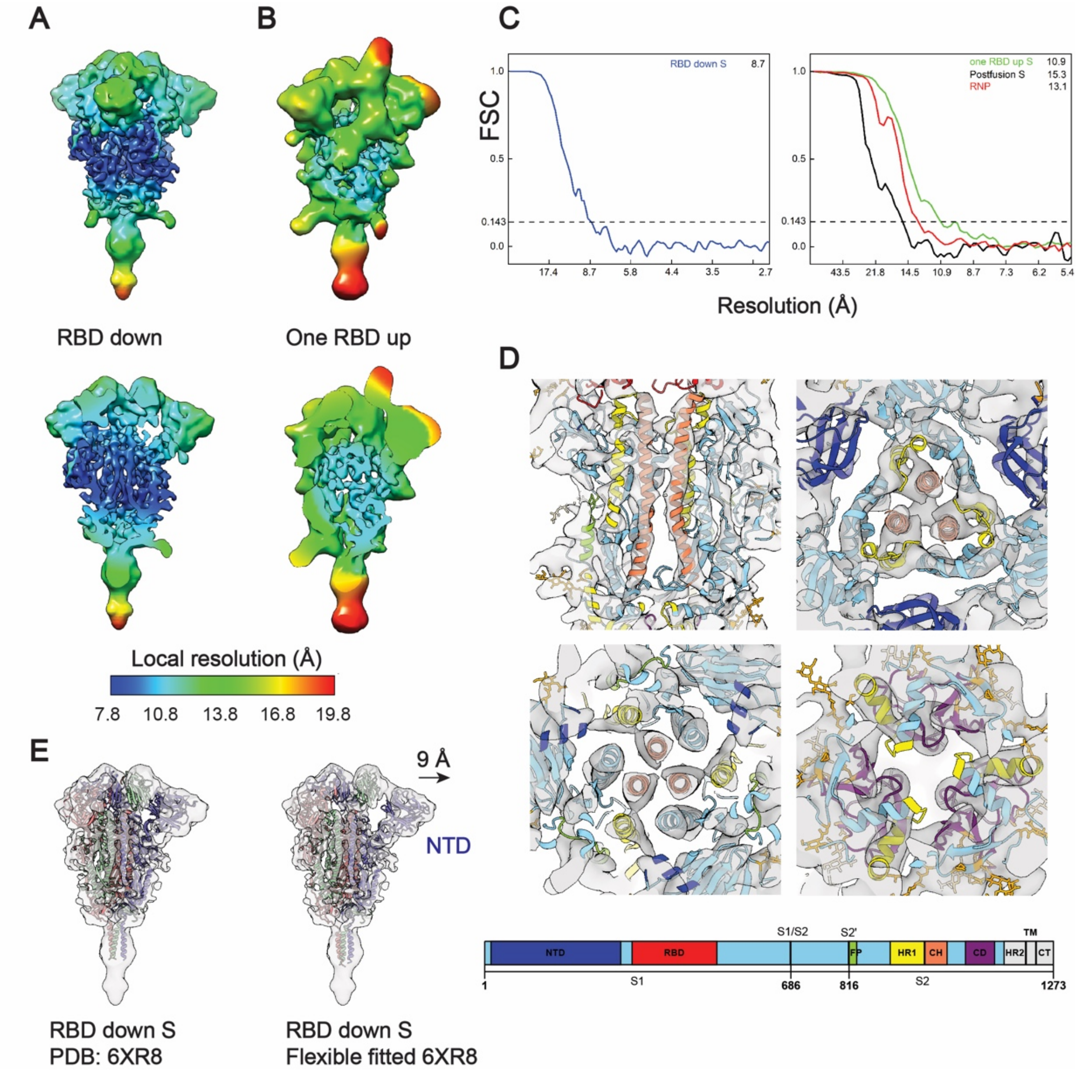


**Figure S3.** Local resolution and Fourier Shell Correlation (FSC) curves. (A, B) Maps of prefusion S in RBD down or one RBD up conformations are colored by their local resolution ranging between 7.8-19.8 Å. Among all domains, the central helical (CH) region is the best resolved, as evidenced by tubular densities of the alpha-helix bundles. (C) Resolution of the spikes in RBD down, one RBD up, the postfusion conformations, and the RNP was estimated from the FSC curves, using a criterion 0.143. (D) The best solved domains of the RBD down S, HR1 and CH from the S2 subunit, are highlighted with the fitted PDB: 6XR8. The domains are colored as in the schematic. NTD, N-terminal domain; RBD, receptor binding domain; S1/S2, S1/S2 cleavage site; S2', S2' cleavage site; FP, fusion peptide; HR1, heptad repeat 1; CH, central helix; CD, connector domain; HR2, heptad repeat 2; TM, transmembrane anchor; CT, cytoplasmic tail. (E) Comparison of the rigidly and flexibly fitted PDB: 6XR8 to the RBD down S. The NTD has shifted by 9 Å away from the CH (centroid distance).

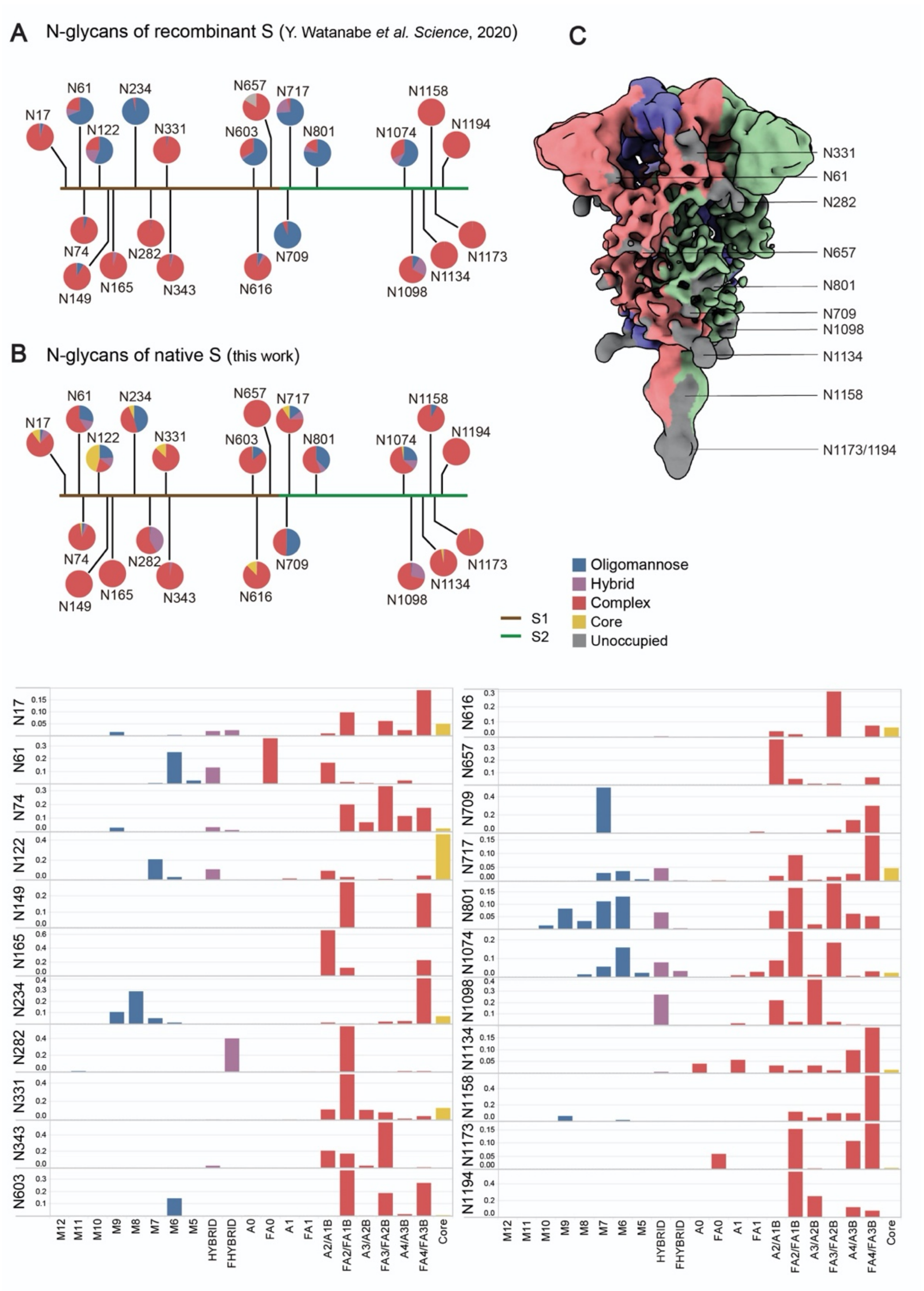


**Figure S4.** N-linked glycans of the native S proteins. (A) Glycan composition of recombinant S, as reported by Y. Watanabe *et al. Science* 2020. (B) Glycan composition of the native full-length S (Oligomannose:M12-M5; Hybrid: Hybrid, Fhybrid; Complex: A0, FA0, A1, FA1, A2/A1B, FA2/FA1B, A3/A2B, FA3/FA2B, A4/A3B, FA4/FA3B). Please check the attached excel data form for the detailed glycan composition. (C) The RBD down S colored by its oligomer subunits. Densities of ten glycans are visible on the map (grey).

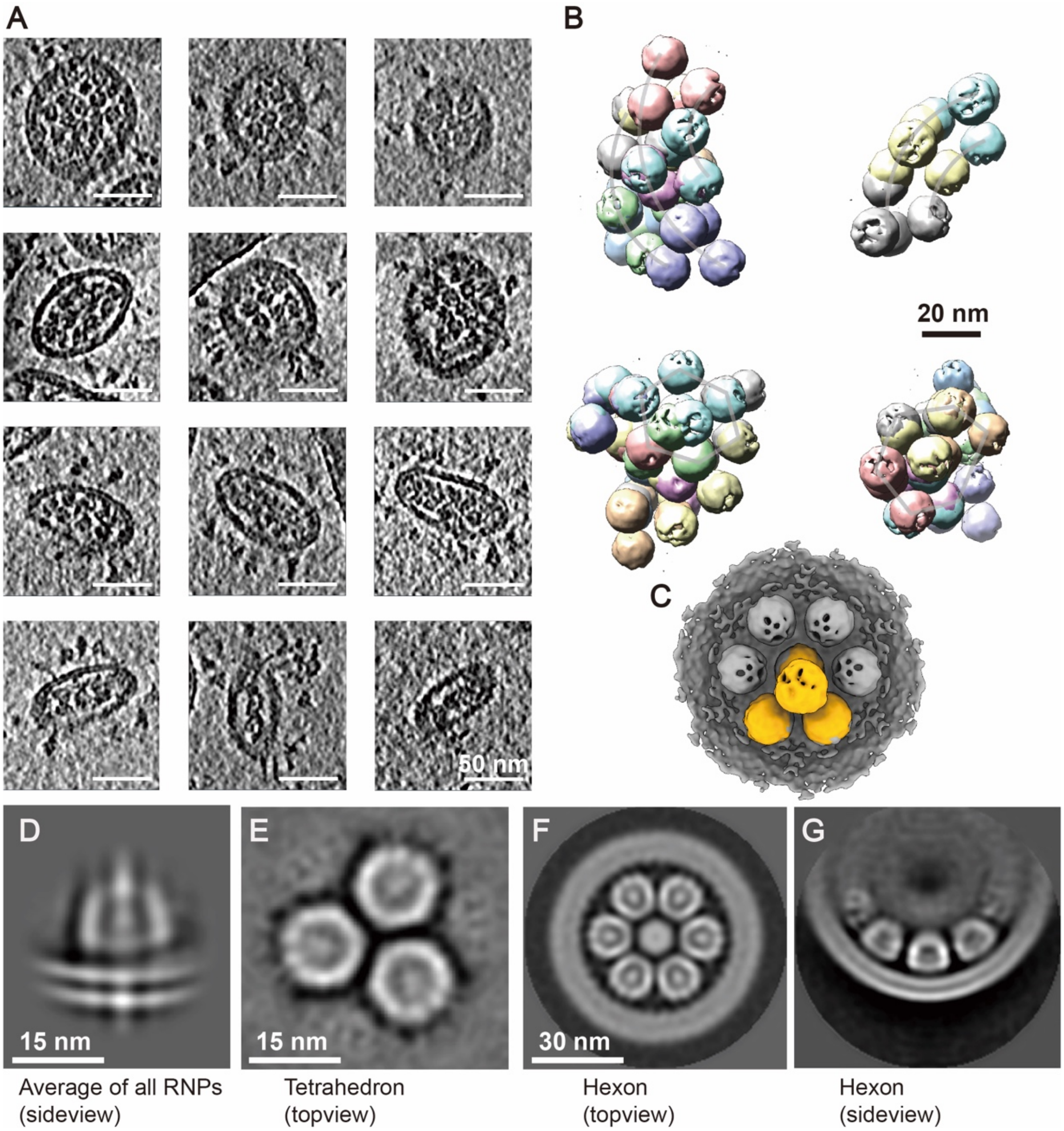


**Figure S5.** Ultrastructure of the RNPs. (A) Example tomogram slices (lowpassed to 80 Å) of *in situ* RNP assembly. Thickness of the slice is 5 Å. (B) RNP tetrahedrons are projected onto their coordinates to show their characteristic higher-order organization. (C) An RNP tetrahedron is overlaid with hexon, showing identical spacing between two neighboring RNPs on either of the assembly. (D) Lipid bilayer density appears, when all RNPs are aligned and averaged using a large spherical mask. (E-G) Topview and sideview of the RNP tetrahedron and hexon reconstructions, showing RNPs are tightly packed in the viral lumen.

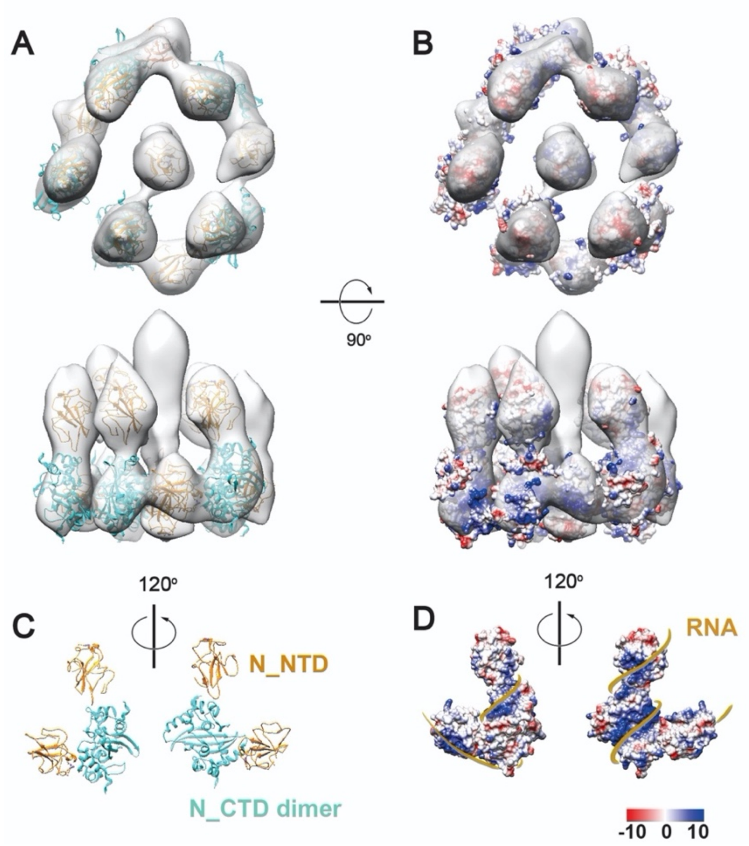


**Figure S6.** A tentative model for the SARS-CoV-2 RNP. (A) Alignment of all single RNPs in regardless of their assembly types using a tight spherical mask revealed a 13.1 Å resolution reverse G-shaped architecture of the RNP, measuring 15 nm in diameter and 16 nm in height. (C) The map was segmented into five head-to-tail reverse L-shaped densities, each fitted with a pair of N (N_NTD: 6WKP, N_CTD: 6WJI) dimerized by the N-CTD, leaving two upper segments unoccupied. Based on the electrostatic potential distribution on the surface of the decamer (B), we propose a tentative model of RNP winded with RNA (D).

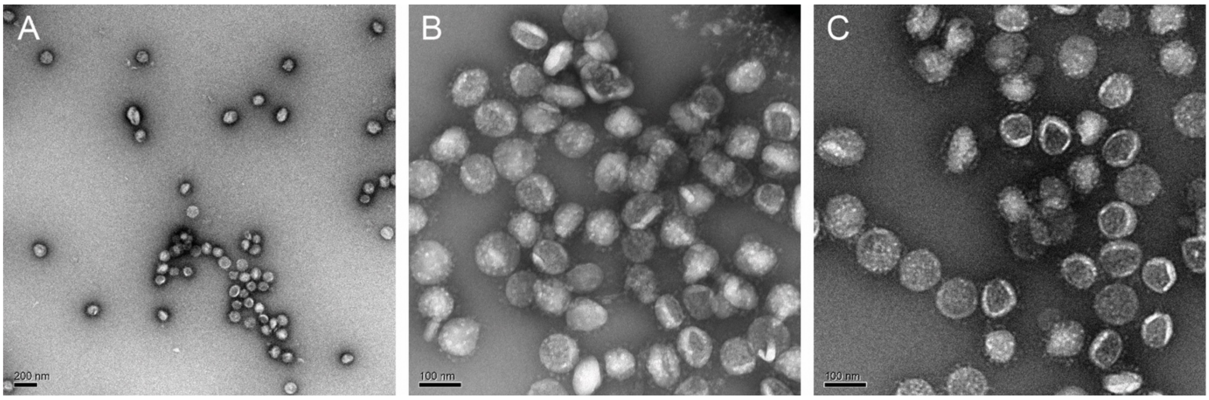


**Figure S7.** SARS-CoV-2 virions remained intact after five cycles of freeze-and-thaw treatment, as shown by negative staining microscopy (A-C).

**Table S1. Cryo-ET Data collection and reconstruction statistics**

| **Data collection** | | | | | | |
|---|---|---|---|---|---|---|
| Microscope | Titan Krios | | | | | |
| Voltage (kV) | 300 | | | | | |
| Detector | Gatan K3 | | | | | |
| Energy filter | Gatan GIF Quantum, 20 eV slit | | | | | |
| Pixel size (Å) | 0.68 (super-resolution) | | | | | |
| Tilt schemes | Dose-symmetric scheme | | | | | |
| Number of tilt-series | 319 | | | | | |
| Number of virions | 2,294 | | | | | |
| Exposure ($e^-$/Å$^2$) | 131.2 | | | | | |
| Defocus range (μm) | -1.7 ~ -5.0 | | | | | |
| Software | SerialEM | | | | | |
| **Reconstruction** | | | | | | |
| Software | Dynamo 1.1.333, Relion 2.0 | | | | | |
| Data set | Prefusion S (RBD down) | Prefusion S (one RBD up) | Postfusion S | RNP (individual) | RNP (tetrahedron) | RNP (hexon) |
| Final number of particles | 25,236 | 21,080 | 1,954 | 18,500 | 3,659 | 2,270 |
| Symmetry imposed | C3 | C1 | C3 | C1 | C3 | C6 |
| Final Resolution (Å) | 8.7 | 10.9 | 15.3 | 13.1 | N/A | N/A |
| Gold-standard | yes | yes | yes | yes | no | no |
| FSC threshold | 0.143 | 0.143 | 0.143 | 0.143 | N/A | N/A |
| Final pixelsize (Å) | 1.36 | 2.72 | 2.72 | 2.72 | 5.44 | 5.44 |
| Map sharpening B-factor (Å$^2$) | -1200 | -2000 | N/A | N/A | N/A | N/A |
